# DySTrack: a modular smart microscopy tool for live tracking of dynamic samples on modern commercial microscopes

**DOI:** 10.64898/2025.12.02.691816

**Authors:** Zimeng Wu, Octavian Voiculescu, Alessandro Mongera, Roberto Mayor, Mie Wong, Jonas Hartmann

**Author notes:** Correspondence: Jonas Hartmann.

## Abstract

Advances in microscopy and bioimage analysis are enabling unprecedented quantitative observation of dynamic biological systems. Smart microscopy closes the loop by feeding back image-derived information to control image acquisition on the fly, paving the way for increasingly autonomous and sophisticated experiments. However, adoption of smart microscopy remains limited primarily to specialists, and even simple tasks such as live tracking of moving samples are still widely handled manually. Here we describe DySTrack, a modular open-source python tool that serves as a minimal bridge between commercial acquisition software and arbitrary image analysis pipelines, allowing users to stick with familiar vendor-developed user interfaces for microscope configuration whilst leveraging the powerful and platform-agnostic python ecosystem for image analysis. DySTrack comes with detailed documentation and with ready-to-use, easy-to-adapt example pipelines that track moving tissues (namely the zebrafish lateral line primordium and the chick Hensen’s node) during embryonic development. We hope DySTrack will contribute to a recent push to make smart microscopy more widely accessible.

## Introduction

Living systems are inherently dynamic, making the ability to visualize, quantify, and manipulate dynamical processes an essential aim of methods development in biology. Light microscopy has long been at the forefront of this endeavor, and recent advances such as array detectors (Huff et al., 2019; Delattre, 2023), adaptive optics (Zhang et al., 2023), new biological sensors (Zhang et al., 2025), optogenetics (Krueger et al., 2019), and computational bioimage analysis (Hallou et al., 2021; Driscoll and Zaritsky, 2021) have upheld a rapid pace of innovation.

A natural and synergistic evolution of these advances is to close the loop between acquisition and analysis by performing on-the-fly image processing and feeding the extracted information back to the microscope to automate and optimize acquisition. This experimental strategy is known as “smart microscopy” (among other names, such as “adaptive-feedback microscopy”) and is thought to have the potential to revolutionize the throughput, quality, reproducibility, and sophistication of bioimaging workflows (Pinkard and Waller, 2022; Carpenter et al., 2023; Morgado et al., 2024; Hinderling et al., 2025a). However, despite the technological feasibility of smart microscopy having been realized decades ago with the introduction of computer-controlled microscopes and having become increasingly viable since, its adoption across research groups remains sparse. Even the relatively basic task of adjusting the field of view to track a moving target is still widely avoided (e.g. by sacrificing resolution, speed, and photon budget for a larger or tiled field of view) or performed through time-consuming manual adjustments that greatly limit throughput.

We concur with a recent assessment (Hinderling et al., 2025a) that this is because smart microscopy has been stuck in a limbo state between sophisticated and bespoke researcher-developed tools on the one hand, and comparably accessible but functionally limited and vendor-locked commercial solutions on the other. Deciding to adopt the former comes with many technical barriers, including the challenge of understanding a bespoke tool deeply enough to convert it to a different use case, but also down-to-earth problems like the simple fact that high-powered microscopes in core facilities do no commonly support open-source control software such as μManager (Edelstein et al., 2010, 2014; Pinkard et al., 2021). For research groups not deeply invested and well-versed in the relevant technical fields, these barriers may appear insurmountable, especially considering ever-looming time constraints. Conversely, the solutions included with commercial microscope control software (such as the Zeiss ZEN Open Application Development^1^ platform or the Nikon NIS-Elements JOBS^2^ module) cannot match the flexibility and rapid advancement of open-source ecosystems such as python. They also lock users into a particular vendor environment and may suddenly lose functionalities when vendor software undergoes major version updates.

We faced this conundrum while looking for a means of autonomously tracking the zebrafish posterior lateral line primordium (pLLP), a highly migratory embryonic tissue (Chitnis et al., 2012; Nechiporuk and Knaut, 2025). In response, we developed a prototype python tool that employs a minimalist and framework-agnostic approach for bridging commercial microscope control software with python-based arbitrary image analysis pipelines, leaving all aspects of microscope configuration to familiar user interfaces whilst simultaneously providing unrestricted access to the power of python-based image analysis. Our prototype proved easy to use, robust, and adaptable to different microscopes and different image analysis workflows. This prompted us to refine it into a generalized, clean, well-tested, and extensively-documented python package that follows modern open-source software conventions, now released under the name DySTrack (“diss track”; Dynamic Sample Tracking).

Here, we describe the architecture and operating principles of DySTrack and showcase its application to three different 3D live tracking problems in two different model organisms (simple center-of mass tracking for drift correction, leading-edge tracking of the collectively migrating zebrafish pLLP, and tracking by model fitting for the regressing chick Hensen’s node) across three different microscope software interfaces (ZEN Black, ZEN Blue, and NIS-Elements). These ready-to-use and easy-to-adapt examples, along with our comprehensive online documentation, will enable users with basic python skills to install, adapt, and use DySTrack for automated sample tracking. Moreover, this work serves as a foundation for ongoing and future efforts to extend the functionality and compatibility of DySTrack to additional microscopes and to more advanced use cases.

## Results

### How DySTrack works

In accordance with its lean and modular design philosophy, DySTrack’s architecture consists of just three components, which are largely independent of each other (Fig. 1a): a microscope macro that controls the microscope’s operations and writes out images, a python image analysis pipeline that reads images and writes out new stage coordinates, and the DySTrack manager, a python tool that mediates between the microscope and the image analysis pipeline.

**Figure 1:**
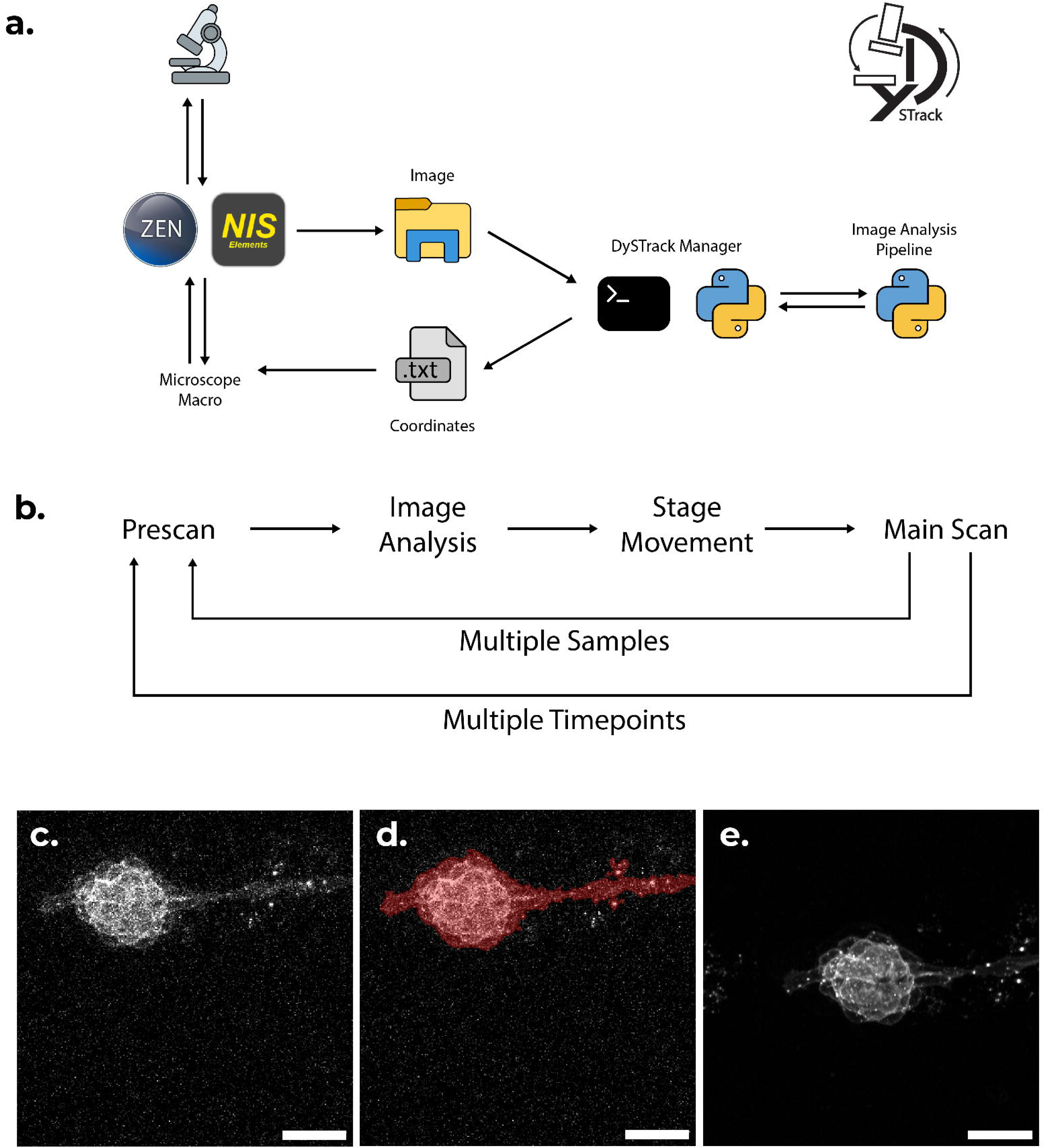
DySTrack architecture and acquisition process. a. Modular architecture of DySTrack, with microscope control handled by vendor software (left), image analysis implemented in python pipelines (right), and the DySTrack manager coordinating the two through a minimal interface (middle). Top right inset: DySTrack logo. b. Acquisition process for a standard experiment where DySTrack adjusts coordinates based on a low-resolution prescan to ensure optimal acquisition of the subsequent high-resolution main scan. c-e. A simple example of DySTrack in action, showing how it re-centers an intentionally heavily offset lateral line neuromast cluster in a transgenic zebrafish embryo. DySTrack reads the prescan (c.), runs an image analysis pipeline that masks the cluster (d.) and finds its center of mass, which is subsequently used for main scan acquisition (e.). All images are maximum z-projections of 3D stacks acquired on the LSM980 in AiryScan2 CO-8Y mode. Fluorescent labeling is krt15:lyn-RFP. Scale bars are 20µm.

Together, these components implement an acquisition loop (Fig. 1b): the microscope moves to a position and acquires an image, the DySTrack manager recognizes the image file, triggers the image analysis pipeline, and feeds the resulting coordinates back to the microscope macro, which corrects the stage position and proceeds with the next acquisition. We usually employ a two-stage process where image analysis is performed on a low-resolution, low-quality, high-speed prescan, which is followed by a high-quality main scan after the stage coordinate update (Fig. 1c-e). This minimizes the time taken by the image analysis pipeline to load and process the image, and maximizes the consistency of the target sample’s location in the main scan field of view. Alternatively, one-step procedures or even more sophisticated multi-step procedures are relatively straightforward to configure (see e.g. the chick Hensen’s node example below, where the main scan is itself a tile scan). In what follows, we describe the functionality and implementation of each component in more detail.

The microscope macro is implemented directly in whichever macro language, automation suite, or API is provided by the microscope vendor, which comes with three advantages. First, users may work with familiar software interfaces when configuring all parameters of the acquisition. Second, the enormous technical complexity of controlling modern microscope hardware remains entirely in the manufacturer’s court, leaving DySTrack users unencumbered by low-level implementation concerns. And third, no custom microscope control software is required, which for many commercial systems (especially confocal laser-scanning microscopes) is either unavailable or would present a significant technical challenge and risk for users/facilities to install. Currently, out-of-the-box support is included for Zeiss’ ZEN Black via the MyPiC pipeline constructor macro (Politi et al., 2018), for Zeiss’ ZEN Blue via an OAD IronPython macro, and for Nikon’s NIS-Elements via the JOBS automation suite. Support for another major vendor is planned for the near future, and our documentation provides detailed instructions for users or vendors to implement support on other systems.

The image analysis pipeline takes the form of an arbitrary python function that is completely independent of the microscope itself, letting users take full advantage of python’s extensive and state-of-the-art image analysis, data analysis, and machine learning ecosystem. A typical pipeline will read a (prescan) image/stack, perform the image analysis steps required to identify the target cell or tissue – for instance by detecting and applying an intensity threshold and morphologically cleaning the resulting foreground mask (Fig. 1d) – and finally extract the (2D or 3D) coordinates of either the center of mass, the leading edge, or another relevant point that should serve as the updated stage position. We showcase a few different approaches in the examples below. It is also possible to use third-party (non-python) software, so long as it either offers a python API or can be executed in a subprocess via a command line interface.

Finally, the DySTrack manager is a python application that awaits new images written out by the microscope, triggers the image analysis pipeline, and returns the coordinates to the microscope macro. Importantly, whilst custom feedback functions can be implemented that communicate directly with the microscope control software (e.g. via an API) or use some other non-standard form of communication (e.g. via the Windows registry for MyPiC), the standard feedback mechanism is to simply write coordinates to a text file. The microscope macro then monitors this text file for new coordinates and triggers stage movement and subsequent acquisitions. This minimal interface is near-universally applicable and has the advantage that microscope software updates will at most require changes to the microscope macro, but not to the DySTrack manager. The manager and image analysis pipelines are therefore well-isolated from microscope-specific considerations. DySTrack manager can be started from the command line or from a Jupyter notebook (or other python process) and it exposes a number of configuration and error handling options that can either be fixed in a configuration file or passed dynamically.

The source code for the DySTrack manager, along with the supported microscope macros and the image analysis pipelines showcased below are available on GitHub^3^ and are extensively documented^4^. We welcome feedback and questions from the community, as well as contributions that add support for other microscope control software, new image analysis pipelines, or other enhancements or bug fixes.

### Tracking migration of the zebrafish lateral line with DySTrack and classical image analysis

The zebrafish pLLP is a placode-derived group of cells that undergo highly coherent, persistent, and directed migration along the lateral midline of the developing embryo’s trunk, assembling and periodically depositing epithelial rosettes that mature into sensory organs, so-called neuromasts (Chitnis et al., 2012; Ghysen and Dambly-Chaudiere, 2007). It is a well-established model system for the study of long-range collective chemotaxis and its dynamic coordination with morphogenesis and cell fate decisions (Haas and Gilmour, 2006; Donà et al., 2013; Durdu et al., 2014; Dalle Nogare and Chitnis, 2017; Wong et al., 2020; Hartmann et al., 2020; Dalle Nogare et al., 2020; Yamaguchi et al., 2022).

Studies of the pLLP rely on high-resolution 3D live confocal fluorescence microscopy to capture the tissue’s rich cellular dynamics, but face the challenge that the pLLP will quickly migrate out of the limited field of view inherent in this imaging modality. This problem is commonly addressed by tiling multiple positions, which limits the time resolution and throughput of experiments, produces large quantities of uninformative image data (i.e. the positions where the primordium is not currently located), and requires subsequent image stitching that may introduce artifacts. In addition, z-drift may occur due to the continued growth of the embryo, due to stage drift, or if embryos are not mounted perfectly horizontally. To reduce the risk of experiment failure, this necessitates preemptively large z-stacks, again wasting time as well as potentially exacerbating photobleaching and phototoxicity. Although many of these challenges are particularly egregious in the pLLP, they are common to long-term high-resolution live imaging of dynamic cells and tissues more generally, making the pLLP an ideal subject for a case study on how to overcome them through automated live tracking by smart microscopy.

To apply DySTrack to this problem, we developed a simple classical image analysis pipeline to mask the pLLP in a low-resolution prescan stack, detect its leading edge, and compute appropriately shifted coordinates for acquisition of the high-resolution main scan (Fig. 2a-c). Briefly, the pipeline performs the following steps: Gaussian smoothing to reduce noise, foreground masking using automated threshold detection with an object-count scheme described in (Hartmann et al., 2020), and removal of all but the largest connected components of the foreground mask. Applied to rapidly and photon-efficiently acquired prescans (Fig. 2a), this method robustly produces a binary mask that captures the entire pLLP (Fig. 2b). By convention, we mount embryos such that the primordium is aligned with the x-axis and migrates from left to right, so the leading edge can be detected simply by finding the right-most voxel of the foreground mask. The new x-coordinate for subsequent acquisition is then calculated such that the leading edge retains a consistent distance from the right border of the field of view. The new y and z coordinates are simply the centroids of the foreground mask. We also include fallbacks in case masking fails or the leading edge has moved out of the field of view in the time since the previous acquisition (see Methods for details).

**Figure 2:**
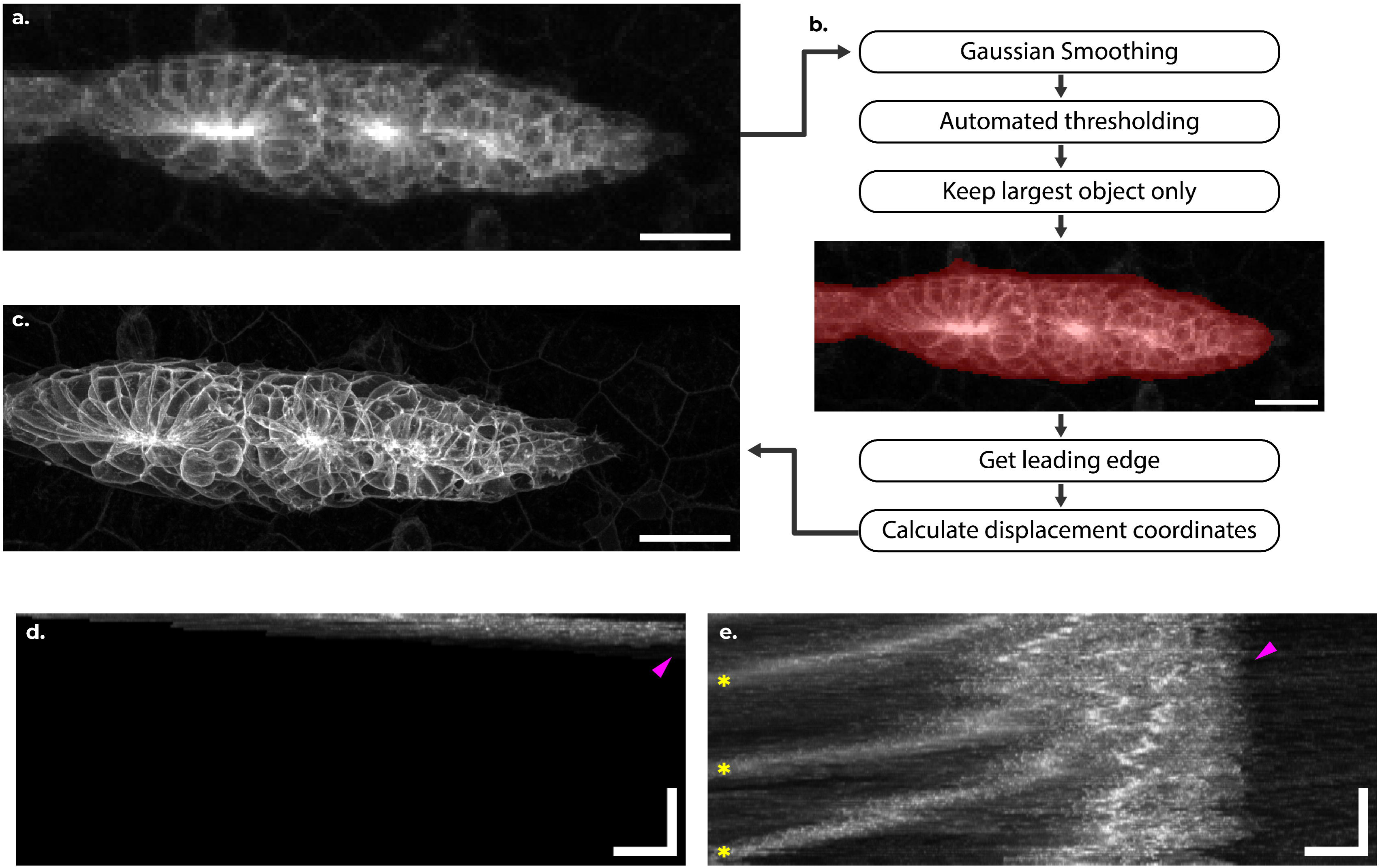
Tracking the zebrafish lateral line primordium. a. Maximum z-projection of a low-resolution prescan of the zebrafish pLLP acquired within seconds (image upsized by a factor of 7.92 to match the main scan below). b. Steps of the classical image analysis pipeline to mask the pLLP (red overlay) and derive new acquisition coordinates. c. Maximum z-projection of high-resolution main scan acquired after adjustment of stage position. Images in a-c are from a DySTrack time course acquired at the Zeiss LSM880 in AiryScan FAST mode; fluorescent labeling is cldnB:lyn-EGFP; scale bars are 20µm; images are cropped in the y-axis for compact presentation (see Methods). d-e. Kymographs of a pLLP tracked over 14.5h. d. Kymograph after deregistration, representing what would have happened with a static field of view. The pLLP leaves this FOV within a short time (magenta arrowhead). e. Kymograph without deregistration, showing robust long-term tracking of the leading edge (magenta arrowhead). Retrograde stripes (yellow stars) are deposited neuromasts. Kymographs in d-e are derived from the top sample in Supp. Movie 2, acquired on the Zeiss LSM980 in AiryScan2 CO-8Y mode; fluorescent labeling is cldnB:lyn-EGFP; horizontal scale bars are 20µm, vertical scale bars are 3.92h.

This simple pipeline, composed of only a handful of basic image analysis operations, enables robust long-term high-resolution live tracking of transgenically labeled lateral line primordia (Fig. 2d-e, Supp. Movie 1) on any microscope supported by DySTrack. Multi-positioning is natively supported (see Fig. 1b), and if desired the resulting positional registration can be reverted in post to recover movies that reflect the true position and speed of the primordium (Supp. Movie 2).

Whilst the pLLP is a particularly extreme case of sample movement, even the largely stationary neuromasts deposited by the primordium may exhibit a degree of drift that can be problematic for long-term time lapse acquisitions. By tracking the center of mass in all dimensions (see Methods for details) as there is no leading edge in this case, DySTrack can be used to stabilize such experiments (see Fig. 1c-e, Supp. Movie 3). This is a basic use case, akin to a “3D software autofocus”, with broad applicability to isotropic tissues in vivo as well as explants and organoids in vitro.

At present, both primordium and neuromast tracking have been extensively tested and successfully used in three different imaging core facilities, across all supported microscopes, with different fluorophores and label intensities, and for several distinct projects (publications forthcoming).

### Tracking regression of the chick Hensen’s node with DySTrack and model fitting

To demonstrate the versatility of DySTrack beyond the pLLP and beyond traditional image analysis pipelines, we implemented tracking of the regressing chick Hensen’s node using a pipeline based on model fitting. Hensen’s node is the amniote gastrulation organizer, which forms at the anterior tip of the primitive streak during gastrulation. As the streak elongates, the node moves anteriorly whilst continuously recruiting cells from the epiblast and producing endoderm underneath, which spreads away from it (Psychoyos and Stern, 1996; Joubin and Stern, 1999). At the onset of elongation, cells stop being added to the node and it regresses caudally, now producing cells for the notochord on the embryo’s midline and the somitic mesoderm flanking it (Voiculescu, 2020; Serrano Nájera and Weijer, 2020), while more posterior portions of the streak continue to act as a gastrulation site (Sheng et al., 2003; Iimura et al., 2007). Thus, the cell populations in and around the node display different types and patterns of movement, and the node moves with respect to all landmarks of the embryo and the substrate on which it grows. In addition, the embryos are large and somewhat variable in the speed of primitive streak elongation and retraction. These characteristics make long-term live imaging of the behavior of cells in and around Hensen’s node particularly challenging – a problem well-suited for DySTrack.

We sought to track the regressing node based on nuclear labeling introduced by electroporation (Voiculescu et al., 2008). The mosaic and variable distribution of labeled nuclei, which exhibits both embryo-to-embryo variation and changes over developmental time, made development of a masking-based pipeline challenging in this case. Furthermore, a simple center-of-mass tracking approach can fail to keep track of the regressing node as clusters of labeled cells leaving the node will “pull” the centroid toward the anterior. We therefore opted to perform a constrained fit of a decreasing sigmoid function to the intensity profile along the anterior-posterior (x) axis, finding that the fitted inflection point robustly tracks the caudal end of the node where signal intensity decreases (Fig. 3a, d). Sticking with this theme, we fitted Gaussian functions to the other axes (y, z) and used the fitted means as new coordinates (Fig. 3b-d). This approach to tracking reflects an alternative way of thinking, wherein one first defines a model of the target (which could be simple functions as in this case, or something more bespoke) and then relies on optimization routines to fit the model to the data, using the resulting parameter values as tracking coordinates. DySTrack’s modular design lets users freely choose and seamlessly deploy such alternative tracking strategies when needed.

**Figure 3:**
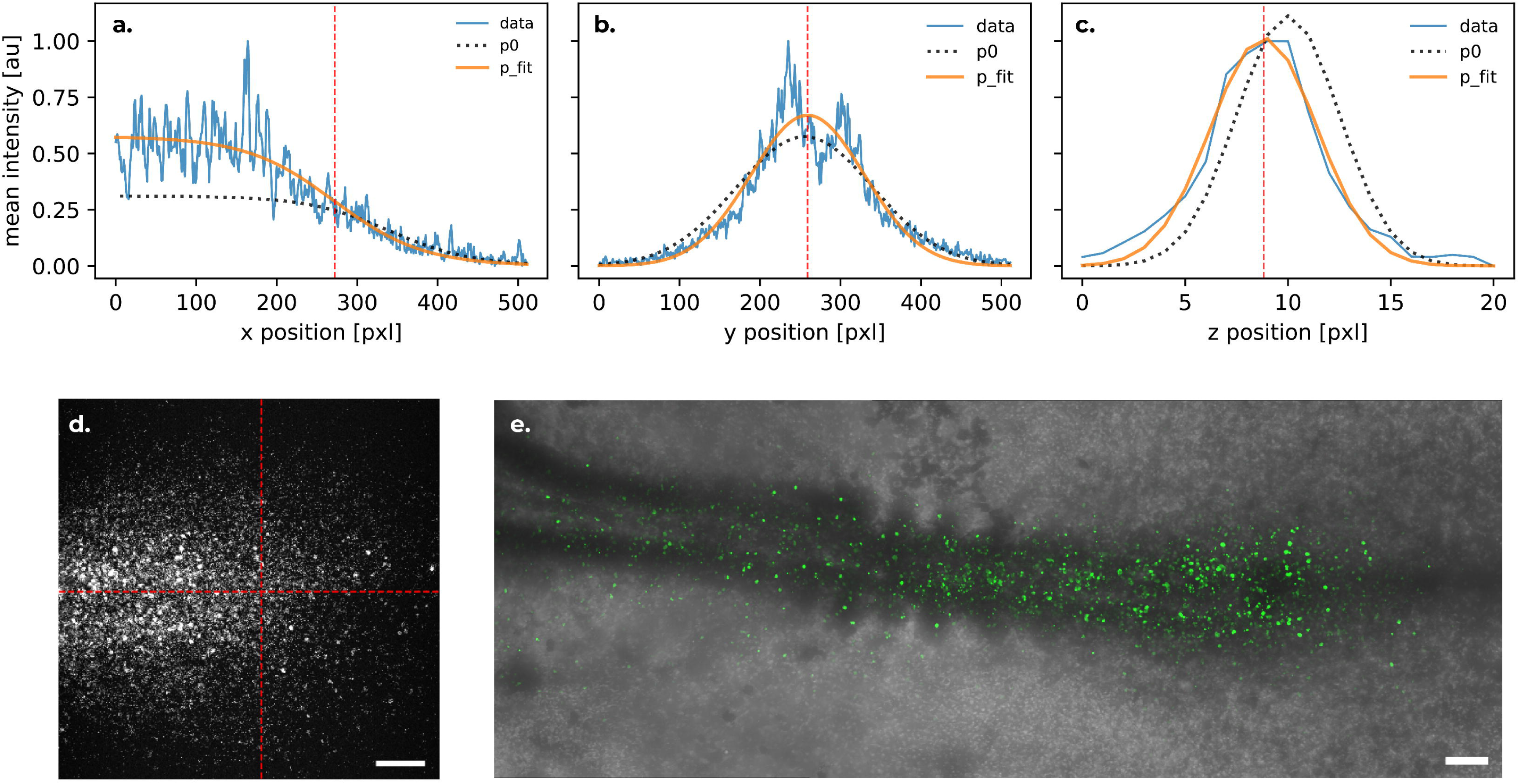
Tracking the chick Hensen’s node. a-c. Intensity profiles (blue lines) of electroporated H2B-EGFP along the axes of a prescan stack, model functions for the initial parameter guess (p0; black dashed lines) and after fitting to data (p_fit; orange lines), and the resulting tracking coordinates (vertical red dashed lines). a. Decreasing sigmoid fit along x-axis. b-c. Gaussian fits along y and z-axes. Note that the fits are relatively close to the initial guesses and the coordinates are close to the image center, as this example is taken from a middle time point (approx. 10h into Supp. Movie 4) where only minor adjustments are required at each step. However, the same pipeline works robustly across the full diversity of node morphologies over time (see Supp. Movie 4). d. Maximum z-projection of the prescan that served as input for a-c, with red dashed lines indicating resulting target coordinates. e. Subsequent main scan acquisition for the same time point, stitched from three tiles, with the rightmost tile being the tracked Hensen’s node. Data was acquired on the Nikon AX R in resonant mode; grayscale is transmission imaging, green is electroporated H2B-EGFP; scale bars are 100µm.

As an additional complication, a single field of view was insufficient in our imaging setup to capture both the node and the cells leaving it. We therefore combined DySTrack with tile-based acquisition in NIS-Elements, using the posterior-most tile (which captures the node itself) for tracking and shifting two additional (more anterior) tiles along with it, resulting in a unique view of chick embryonic axis development (Fig. 3e, Supp. Movies 4, 5).

## Discussion

We described DySTrack, a python-based smart microscopy tool for dynamic sample tracking in 3D on commercial microscopes available in most facilities. DySTrack talks to microscope software through a minimalist interface to maximize modularity, leaving all microscope control and configuration to manufacturer software and simultaneously providing unlimited access to the python ecosystem for image analysis. We demonstrated how DySTrack changes the game for high-quality time-lapse imaging of highly migratory tissues such as the zebrafish pLLP, and that it can readily be adopted to different tracking problems, such as chick node regression. DySTrack is available as an open-source python package that includes an automated test suite and extensive documentation. We expect that its continued development by ourselves and its user community will extend its applicability to different use cases and microscopes.

Modern methods of generating static, single-time-point biological data have been scaled up to such high levels of throughput and automation that data analysis and interpretation – not data acquisition – are now their main bottleneck. However, understanding inherently dynamic systems from static data remains a fundamental obstacle. Live time-lapse microscopy resolves this problem but has not seen the same gains in throughput, as it faces multiple trade-offs, struggles with high rates of failed experiments, and often requires laborious manual supervision. Smart microscopy could potentially alleviate all of these drawbacks, yet its adoption thus far remains restricted to specialist labs and advanced applications (Hinderling et al., 2025a).

To accelerate adoption, low-barrier entry points are needed for biologists and facilities to explore the power of smart microscopy without a substantial up-front time investment to develop and test bespoke solutions. Recently, several new smart microscopy tools have been released that each have a somewhat different focus and design philosophy, but ultimately all aim to make smart microscopy more widely accessible. Examples include general smart microscopy toolkits such as AutoScanJ (Tosi et al., 2021) and MicroMator (Fox et al., 2022), frameworks for event-driven microscopy and advanced HCS applications such as CelFDrive (Brooks et al., 2024), EDA (Mahecic et al., 2022), and DDM (André et al., 2023), and tools for optogenetic control such as pyCLM (Oatman et al., 2025), Outcome-Driven Microscopy (Passmore et al., 2024), and rtm-pymmcore (Hinderling et al., 2025b). DySTrack joins the ranks of these tools as an accessible entry point for dynamic sample tracking with low adoption cost and complexity.

Collectively, this growing ecosystem affords researchers the freedom to choose a tool that suits their experimental needs, prior expertise, and existing infrastructure. We hope this will accelerate adoption and in turn stimulate further innovation and standardization (Hinderling et al., 2025a), resulting in a positive feedback loop that propels smart microscopy from infancy to maturity.

## Materials and Methods

### Zebrafish handling and husbandry

Zebrafish husbandry and experiments were conducted according to UCL Fish Facility standard protocols and under Home Office project license PP0482739 awarded to MW, according to the UK Animal Scientific Procedures Act (1986). Embryos were kept in Petri dishes in fish water (51mM NaCl, 0.171mM KCl, 0.331mM CaCl2, 0.331mM MgSO4) in an incubator at 28-30°C without light cycle, and were staged as previously described (Kimmel et al., 1995).

The following published transgenic lines were used: krt15:lyn-RFP (Wong et al., 2020) outcrossed to WT in Fig. 1c-e, cldnB:lyn-EGFP (Haas and Gilmour, 2006) outcrossed to WT in Fig. 2 and Movies 1 and 2, and krt15:lyn-RFP; cxcr7b:cxcr7b-EGFP (Wong et al., 2020) outcrossed to WT in Movie 3.

### Zebrafish lateral line imaging

Pigmentation of embryos was prevented by treating them with 3mg/mL N-phenylthiourea (PTU) (Sigma) starting at 25hpf. Embryos at 30-36hpf were manually dechorionated with forceps, anaesthetized with Tricaine, screened at the stereoscope for transgenic marker expression, and embedded in 0.8% low melting agarose (Sigma) in fish water containing 0.016% Tricaine. Embryos were mounted on their side in a 35mm MatTek glass-bottom dish (coverslip thickness 0.17mm). Once the agarose solidified, fish water containing 0.016% Tricaine was added to the dish. The incubation temperature throughout imaging was kept constant at 28°C.

Imaging was performed on an inverted Zeiss LSM880 microscope (Figs. 2a-c and Movie 1) and on an inverted Zeiss LSM980 microscope (Figs. 1c-e, 2d-e and Movies 2, 3). Both setups used a Zeiss 40X 1.2NA water immersion objective. Acquisition speed was maximized using a piezo z-drive, bi-directional scanning, and no averaging. Main scans were performed in AiryScan FAST mode (LSM880) or AiryScan2 CO-8Y mode (LSM980), with high pixel densities and optimal z-sectioning (Fig. 1e: 0.114µm in xy and 0.230µm in z; Fig. 2c and Movie 1: 0.103µm in xy and 0.225µm in z; Fig. 2d-e and Movie 2: 0.099µm in xy and 0.210µm in z; Movie 3: 0.099µm in xy and 0.210µm in z). Prescans were acquired in confocal mode (1AU pinhole), with reduced laser power and increased gain, and with greatly reduced pixel density and z-resolution (Fig. 1c-d: 0.471µm in xy and 2.000µm in z; Fig. 2a-b and prescans for tracking Movie 1: 0.817µm in xy and 2.932µm in z; prescans for tracking Fig. 2d-e and Movie 2: 0.420µm in xy and 2.000µm in z; prescans for tracking Movie 3: 0.530µm in xy and 2.000µm in z). Z-stack height was chosen such that the samples were fully captured with approx. 15% extra space on either side in main scans and approx. 25% extra in prescans. AiryScan processing was performed using the respective microscopes’ built-in tools with “auto” and “3D processing” settings selected. Images in Fig. 2a-c were cropped in the y-axis by 120px at the top and 139px at the bottom, for compact presentation. The kymographs in Fig. 2d-e were generated from a rectangular stripe spanning the full x-axis and reaching from pixel 281 to pixel 348 in y (67px height).

### Chick embryo handling and electroporation

Fertilized hen’s eggs (Charles Rivers) were incubated at 38°C to reach stage Hamburger and Hamilton stage HH3+ (Hamburger and Hamilton, 1951). They were then electroporated with a construct driving the ubiquitous expression of H2B-EGFP and cultivated in New culture for 8 hours following the protocol described in (Voiculescu et al., 2008).

### Chick Hensen’s node imaging

The glass rings carrying the embryos selected for imaging were then transferred into customized, 3D-printed plastic dishes tailored similar to those described previously (Voiculescu and Stern, 2012), in order to allow imaging through the endoderm. Briefly, the sealed dishes were designed to encase the glass ring keeping the vitelline membrane under appropriate tension and ensure normal development of the embryo, and the culture wells described previously were raised to 3mm and filled with albumen containing 1.2% agar-agar so that the embryo is raised to within 1mm of the glass coverslip covering the glass ring and allowing dry lenses of relatively long working distance to be used. The temperature of the microscope hood was maintained at 38°C during imaging, with no need for humidification.

Data shown in Fig. 3 and Movies 4, 5 were acquired on a Nikon AX R NSPARC inverted point scanning confocal microscope in AX resonant mode, with a Nikon 20X 0.45NA (8.2mm working distance) dry objective, using a piezo z-drive, no averaging, and pinhole set to 1.0 AU. DySTrack main scans were performed with high pixel densities and relatively narrow z-sectioning (Fig. 3e and movies 4, 5: 0.863µm in xy and 3.000µm in z), whereas the pixel density and z-resolution of prescans was greatly reduced (Fig. 3a-d and prescans for tracking movies 4, 5: 1.726µm in xy and 10.0µm in z) and gain was increased. Z-stack height was chosen such that the node region of the sample was fully captured, with again as much extra space on either side in main scans, and about 1.5x as much space on either side in prescans (the relatively large amount of extra space was included to anticipate the slight angling/bending embryos exhibited at later stages). For main scans, two additional tile positions anterior of the tracked node position were added with 15% overlap in x and matched z-coordinates. The acquired main scan tiles were individually denoised in NIS-Elements (Nikon) using GA3 Denoising with default settings, then maximum z-projected and subsequently stitched in ImageJ using the Grid/Collection Stitching plugin (Preibisch et al., 2009) with default settings.

### Zebrafish neuromast center-of-mass tracking pipeline

Prescans were smoothed with a 3D Gaussian filter (σ=3) to reduce noise. The threshold_otsu function from scikit-image was applied to detect foreground pixels. Only the largest connected component was retained and its center of mass in all three dimensions was returned as the new acquisition coordinates for DySTrack. In z, a safety limit was imposed that restricts stage movement to at most 1/10th of the depth of the z-stack per time point. The DySTrack center-of-mass tracking pipeline also supports masking with object-count thresholding (see below) or direct calculation of the center of mass from input pixel intensities (without masking).

### Zebrafish pLLP leading edge tracking pipeline

Prescans were smoothed with a 3D Gaussian filter (σ=3) to reduce noise. Object-count thresholding (Hartmann et al., 2020) was applied to detect foreground pixels. Briefly, this works by applying a series of thresholds (here one for each intensity value in the 8bit range) and counting the number of objects (connected components) in each mask. The resulting distribution of object counts over threshold values is smoothed using a 1D Gaussian filter (σ=3). In this profile, low thresholds will be associated with high object counts because background noise is captured as small objects. As the threshold increases, the count will suddenly drop dramatically and then remain relatively stable or increase slightly as foreground objects start to be split. The optimal threshold thus lies just after the sudden decrease in the profile, which is detected based on the following criteria: it must be greater than the threshold value at the peak of the object-count distribution, and its associated object count must either be lower than the peak count by at least a factor of 2 or must be a local minimum, and there must be at least one object present at that threshold (otherwise the next lower threshold with >0 objects is chosen as a fallback). Although somewhat lacking in elegance, this algorithm has proven extremely robust.

Once the optimal threshold was found and applied, only the largest connected component was retained as the final mask for the pLLP, and its center of mass was used as the new DySTrack acquisition coordinates in the z and y dimensions. In z, a safety limit was imposed that restricts stage movement to at most 1/10th of the depth of the z-stack per time point. In the x dimension, the leading edge was detected as the frontal-most voxel within the mask and the new x-coordinate was computed such that the subsequent acquisition would place the leading edge 1/5th of the image size away from right-hand border. Two fallbacks were put in place to handle occasional issues. First, if the mask touches the right-hand border of the prescan, the pLLP is assumed to have moved too far between acquisition, and the field of view is moved rightward by a “catch-up distance” of 1/5th of the image size (note that this differs from the standard case where much less movement may be required to place the leading edge 1/5th away from the border). Second, if the leading edge is detected behind the midpoint of the image, this is assumed to be due to a masking error resulting from unusually strong variation in the brightness of the pLLP’s labeling along the x-axis. In this case, the field of view is moved rightward by a default distance of 1/8th of the image size. These fallbacks are hardly ever triggered when tracking bright and uniformly-labeled primordia, but on the rare occasions where they come into play, they may rescue the entire time lapse for a sample that might otherwise be lost.

### Chick Hensen’s node tracking pipeline

For performance, input prescans were converted from 16bit to 8bit by linear rescaling of the range between their minimum and maximum value to the range between zero and 255. A Gaussian probability density function was defined to model the intensity distribution along the z and y axes:

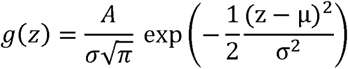

Substitute *y* for *z* for the y-axis case. The intensity profiles along z and y were computed as the mean intensities over both other dimensions, and were again linearly rescaled, this time into floating-point values between zero and one. The Gaussian was then fit to each of these profiles using the curve_fit function from scipy.optimize, with the following initial guesses for parameters: *A* is the sum of the respective profile, µ is the center of the respective axis, and σ is 1/8th of the extent of the z-axis or 1/6th of the extent of the y-axis, respectively. The fitted values for µ were returned to DySTrack as the new z and y coordinates. In z, a safety limit was imposed that restricts stage movement to at most 1/5th of the depth of the z-stack per time point.

A decreasing sigmoid function was defined to model the posterior boundary of Hensen’s node along the x-axis:

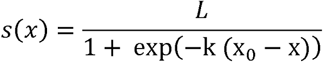

The intensity profile along x was computed as the mean intensities over both other dimensions, but in y including only values within a stripe of width 2σ*_y_* centered on the detected coordinate µ_y_ (using the values from the Gaussian fit above) to focus on the embryo’s midline where the node’s signal is most dense. This profile was again linearly rescaled to floating-point values between zero and one. The sigmoid was then fit using curve_fit with these initial guesses for parameters: *L* is the mean of profile intensities, k scales the steepness of the curve with initial guess 10/X (where X is the extent of the x-axis), and x_O_ is the inflection point with initial guess 2X/3. For additional robustness, bounds were imposed on the parameters: *L* must lie between the minimum value and 1.2 times the maximum value of the profile, *k* must lie between 1/X and 100/X, and x_O_ must lie between X/6.0 and X/1.2. All initial guesses and bounds were chosen to ensure maximum robustness based on data from pilot experiments. The fitted value for x_O_ was returned to DySTrack as the new x coordinate.

### Software

Version indicators provided in this section reflect software supported or depended upon as of DySTrack version 1.0.0, released December 2025 and archived at Zenodo^5^. DySTrack is built with python (tested under versions 3.10 through 3.13), using git and GitHub for version control, miniforge for package management, pytest for automated testing, and sphinx for documentation. Notable scientific python dependencies include numpy ≥1.24 (Harris et al., 2020), scipy ≥1.10 (Virtanen et al., 2020), matplotlib ≥3.7 (Hunter, 2007), and scikit-image ≥0.24.0 (van der Walt et al., 2014). Microscope images of various formats are opened with bioio ≥3.0.0 (Brown et al., 2023) with appropriate plugins. Jupyter notebook <7 (Granger and Pérez, 2021) was used for prototyping and to implement the deregistration workflow. Some image visualization and processing was performed with Fiji 1.54p (Schindelin et al., 2012). The Zeiss LSM880 was controlled with ZEN Black 2.3 SP1, the Zeiss LSM980 with ZEN Blue 3.12, and the Nikon AX R with NIS-Elements ER 6.10.01. Macro control for ZEN Black is implemented with MyPiC commit 6682255 (Politi et al., 2018). Supported operating systems for DySTrack are Microsoft Windows 10 and 11.

## Author Contributions

JH developed the initial prototype. ZW and JH developed and tested the greatly improved modern version presented here, with support by MW and RM. JH and OV developed the application to chick node tracking with support by AM. ZW and JH wrote the manuscript. All authors reviewed and edited the manuscript.

## Acknowledgments

We thank the UCL Centre for Cell & Molecular Dynamics (CCMD) facility for support with microscopy, especially Virginia Silio, Michael Redd, and Alan Greig. We thank the UCL zebrafish facility for zebrafish stock maintenance. For support with integration development, we thank Nicolas Sergent (Zeiss) for ZEN Blue and Robert Tetley (Nikon) for NIS-Elements. JH further thanks Darren Gilmour, Antonio Politi, Elisa Gallo, Sabine Görgens, and the Advanced Light Microscopy Facility (ALMF) at EMBL Heidelberg, especially Aliaksandr Halavatyi and Christian Tischer, for support during prototype development.

## Data and resource availability

The code for DySTrack is freely available (under the MIT license) and will be openly maintained on GitHub: https://github.com/WhoIsJack/DySTrack. The version described here is 1.0.0 and is archived on Zenodo: https://zenodo.org/records/17782770. The repository includes prescan data necessary for testing DySTrack’s functionalities. It also includes the raw files for the associated documentation, which is hosted online here: https://whoisjack.github.io/DySTrack. Raw rata underpinning the figures and movies in this publication are available from the corresponding author, JH, upon reasonable request.

## Supplementary movie legends

### Supplementary Movie 1: High-resolution tracked lateral line primordium

Maximum z-projected high-resolution (main scan) time course of a transgenically labeled pLLP (green; cldnB:lyn-EGFP) tracked with DySTrack on a Zeiss LSM880 in AiryScan FAST mode (40X 1.2NA water objective). Scale bar 20µm, time resolution 5min, and total time course duration 8.33h. Movie is JPEG compressed.

### Supplementary Movie 2: Multi-position pLLP acquisition with subsequent deregistration

Maximum z-projections of three simultaneously tracked lateral line primordia (green; cldnB:lyn-EGFP). The live registration applied by DySTrack was reverted in post to reconstruct the samples’ actual movement in space (but note that positions in the y axis are not reflective of embryo locations in the dish). Raw data were acquired on a Zeiss LSM980 in AiryScan2 CO-8Y mode (40X 1.2NA water objective). Scale bar is 50µm, time resolution 10min, and total time course duration 14.66h. To cope with the size of this reconstruction, images were downsampled by 2x with bicubic interpolation, and the movie is JPEG compressed.

### Supplementary Movie 3: Long-term high-resolution tracking of a deposited neuromast

Maximum z-projections of a main scan time course of a deposited zebrafish neuromast, whose center of mass was tracked with DySTrack (right panel) based on the transgenic membrane label krt15:lyn-RFP (yellow). A second channel not used for tracking was also imaged, showing a transgenically labeled atypical chemokine receptor (cxcr7b:cxcr7b-EGFP; magenta). The movie was acquired on the Zeiss LSM980 in AiryScan2 CO-8Y mode (40X 1.2NA water objective). The live registration applied by DySTrack was reverted in post to reconstruct the sample’s actual movement in space (de-registered stack; left panel) with the trace of DySTrack centroid positions indicated with white/cyan squares, revealing the sample drift that DySTrack has corrected. Scale bar is 20µm, time resolution 15min, and total time course duration 12h. Movie is JPEG compressed.

### Supplementary Movie 4: Multi-tile acquisition of tracked Hensen’s node

Maximum z-projected main scan time course of the chick Hensen’s node tracked with DySTrack on a Nikon AX R in resonant mode (20X 0.45NA 8.2mm-WD dry objective). Grayscale is transmission imaging, green is electroporated H2B-EGFP. The entire view is composed of three stitched acquisitions (15% overlap) with the right-most field of view being live tracked. The movie shows the node forming and regressing to the right (posterior) as somites are laid down. Note that the embryo begins to curve out of the plane in the second half of the movie, yet despite this issue – and despite changes in the morphology and labeling of the node – the model-based DySTrack pipeline is able to maintain stable tracking. Scale bar is 200µm, time resolution 10min, and total time course duration 22h. Movie is JPEG compressed.

### Supplementary Movie 5: Deregistered tracked Hensen’s node

The same data as shown in Supp. Movie 4, but deregistered in post to visualize the actual spatial displacement of the node during regression. Scale bar is 100µm, time resolution 10min, and total time course duration 22h. To cope with the size of this reconstruction, images were downsampled by 2x with bicubic interpolation, and the movie is JPEG compressed.

## Footnotes

1 https://github.com/zeiss-microscopy/OAD

2 https://www.microscope.healthcare.nikon.com/en_EU/products/software/nis-elements/nis-elements-jobs

3 https://github.com/WhoIsJack/DySTrack

4 https://whoisjack.github.io/DySTrack

5 https://zenodo.org/records/17782770

